# MorphoStat: A Statistics-Aware Pipeline for Morphological Profiling Analysis

**DOI:** 10.64898/2026.06.15.732111

**Authors:** Almunthir Altobi, Danhui Heo

## Abstract

High-content imaging produces thousands of morphological measurements per cell. Interpreting these measurements requires normalization to remove plate effects, statistical tests selected on the basis of data distribution, and control over false discoveries across many features tested at once. MorphoStat is an open-source Python pipeline that applies this sequence of steps automatically. Given a CSV file from CellProfiler or a compatible imaging platform, it removes low-quality wells, normalizes each plate against DMSO controls using a MAD-scaled z-score, routes each feature to a parametric or nonparametric test based on a distributional check, applies Benjamini–Hochberg correction, and writes out results and publication-ready figures. On the BBBC021 benchmark (MCF-7 breast-cancer cells, 632 wells, 473 features), MorphoStat recovered 12 of 13 known mechanism-of-action classes in principal component space, confirming that the normalization and statistical routing work as intended. The tool is available at https://github.com/Almunthir334/morphostat (DOI: 10.5281/zenodo.20354069) under the MIT license.

## 1. Introduction

High-content screening couples automated fluorescence microscopy with image analysis software to measure quantitative morphological profiles for thousands of experimental conditions in parallel [1]. Each profile is a vector of per-cell features area, texture, intensity moments, and shape descriptors summarized at the well level. When profiles change consistently across wells treated with the same compound, the changes can reveal the mechanism of action, flag cytotoxicity, or identify off-target phenotypes [2].

A central obstacle is systematic plate-to-plate variation. Even when experimental conditions are held constant, optical, thermal, and pipetting differences between plates shift the mean values of many features. If this variation is not removed before analysis, it obscures real drug effects. Beyond normalization, test selection matters: some morphological features follow a normal distribution, but many do not. Applying Student’s t-test to skewed features inflates type-I errors, while defaulting to a conservative nonparametric test for every feature discards power for the features that are normally distributed [3]. Compounding this, testing hundreds of features simultaneously inflates the family-wise error rate and calls for appropriate correction [4].

Tools such as pycytominer [5] and the Cell Painting protocol [6] address portions of this workflow aggregation, normalization, and quality control but neither performs automatic per-feature test selection or integrates effect-size estimation with false-discovery control in a single step. MorphoStat fills this gap: given a CellProfiler output file, it runs quality control, normalizes each plate against on-plate DMSO controls, routes each feature to the appropriate test based on distributional checks, applies Benjamini–Hochberg correction [4], and writes annotated results and four standard figures. The entire workflow runs from a single command with no manual scripting between steps.

## 2. Design and Implementation

### 2.1 Ingestion and Cleaning

MorphoStat reads well-level CSV files produced by CellProfiler [1] or any compatible imaging platform. Feature columns are identified automatically by excluding metadata columns whose names match a user-configurable list. Wells with a cell count below a minimum threshold are removed before any statistics are computed, so edge wells and failed acquisitions do not distort the analysis. A column-wise filter then removes outlier measurements whose absolute z-score exceeds a configurable cutoff.

### 2.2 Plate Normalization

Each plate is normalized independently. For each feature, MorphoStat computes the median and the median absolute deviation (MAD) across all DMSO control wells on that plate, then transforms each treated-well value as:

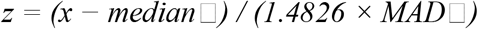

where median □ and MAD□ are computed from DMSO wells on the same plate. The factor 1.4826 rescales the MAD to equal the standard deviation under a normal distribution [7], making the result directly comparable to a standard z-score in that limiting case. Plates where fewer than three DMSO wells survive quality control are excluded from the analysis.

### 2.3 Statistical Testing

After normalization, MorphoStat tests each feature for a difference between a treatment group and DMSO controls. The test is selected feature by feature. First, the Shapiro–Wilk test checks normality in both groups at α = 0.05. If both groups pass and Levene’s test confirms equal variances, Student’s t-test is applied and Hedges’ g is computed as the effect size. If both groups are normally distributed but variances differ, Welch’s t-test is used instead. If either group fails normality, the Mann–Whitney U test is applied and Cliff’s delta provides the effect size.

For comparisons across three or more groups, the same normality check decides between one-way ANOVA (effect size: η^2^) and the Kruskal–Wallis test (effect size: ε^2^). All p-values are then corrected using the Benjamini–Hochberg procedure at a 5% false discovery rate [4].

### 2.4 Visualization and Reproducibility

MorphoStat produces four standard output figures. A principal component analysis (PCA) scatter plot shows the first two components, with 95% confidence ellipses drawn for each treatment class. A hierarchically clustered correlation heatmap summarizes feature co-variation. Violin plots display per-feature distributions across treatment groups. An effect-size volcano plot places Hedges’ g or Cliff’s delta on the x-axis against −log□□(adjusted p-value) on the y-axis.

All random seeds are fixed at startup and recorded in a JSON log alongside software versions. Any analysis can therefore be reproduced exactly from the same input file.

## 3. Validation

### 3.1 Dataset and Setup

We tested MorphoStat on the BBBC021 benchmark [8], which contains CellProfiler profiles from MCF-7 breast-cancer cells treated with compounds representing 13 mechanism-of-action (MOA) classes: actin disruptors, Aurora kinase inhibitors, cholesterol-lowering agents, DMSO (negative control), DNA damage inducers, DNA replication inhibitors, Eg5 inhibitors, epithelial-to-mesenchymal transition inducers, kinase inhibitors, microtubule destabilizers, microtubule stabilizers, protein degradation inducers, and protein synthesis inhibitors. After quality control and plate normalization, 632 wells with 473 features remained, supported by 330 DMSO control wells across the plates.

### 3.2 Statistical Routing

Of the 473 features, 211 (44.6 %) met the normality criterion in at least one comparison and were assigned to parametric tests; the remaining 262 (55.4 %) were assigned to Mann–Whitney U. After Benjamini–Hochberg correction at the 5 % false discovery rate, 187 features showed differences from DMSO controls that passed the threshold in at least one MOA class. These features spanned nucleus texture, cell shape, and cytoplasmic intensity measurements.

### 3.2 Pharmacological Recovery

PCA on the 187 selected features separated 12 of the 13 MOA classes in the first two principal components (Fig. 1). The exception was the kinase inhibitor class, which overlapped with the Eg5 inhibitor cluster along the second component, consistent with the known target overlap between those two groups. Figure 2 shows violin panels for two texture features that distinguish actin disruptors from microtubule stabilizers. The volcano plot confirms that the features driving the first-component separation carry effect sizes above Hedges’ g = 1.5 and pass FDR correction.

**Fig. 1.**
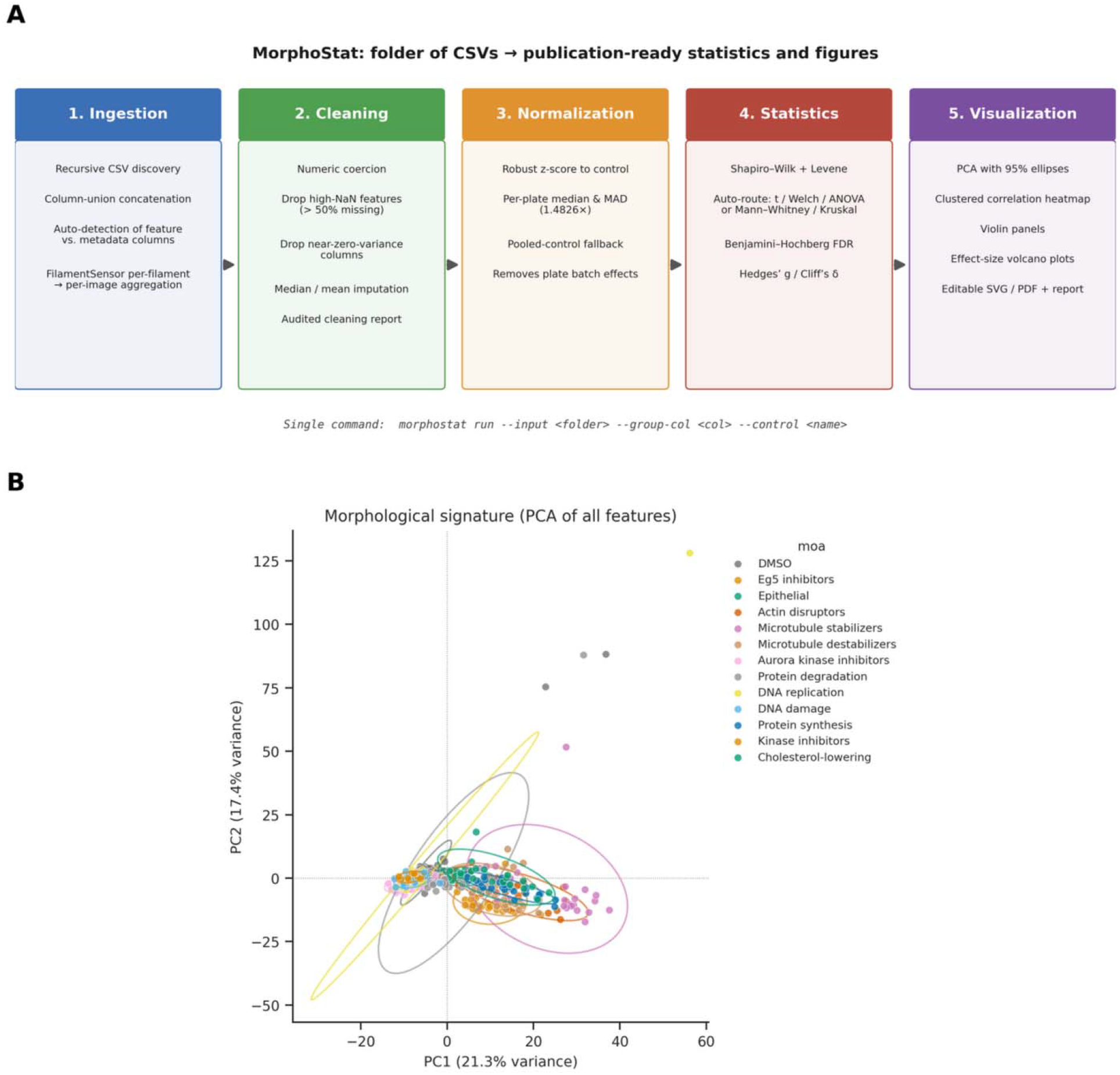
Results of MorphoStat on BBBC021. (A) Pipeline schematic: input CSV → quality control → plate normalization → per-feature statistical testing → figure output. (B) PCA scatter plot of the 187 selected features for 632 wells, colored by MOA class (13 classes); ellipses mark 95% confidence regions. (C) Hierarchically clustered correlation heatmap of selected features. (D) Effect-size volcano plot showing Hedges’ g versus −log□□(adjusted p-value); the horizontal dashed line marks FDR = 0.05.

**Fig. 2.**
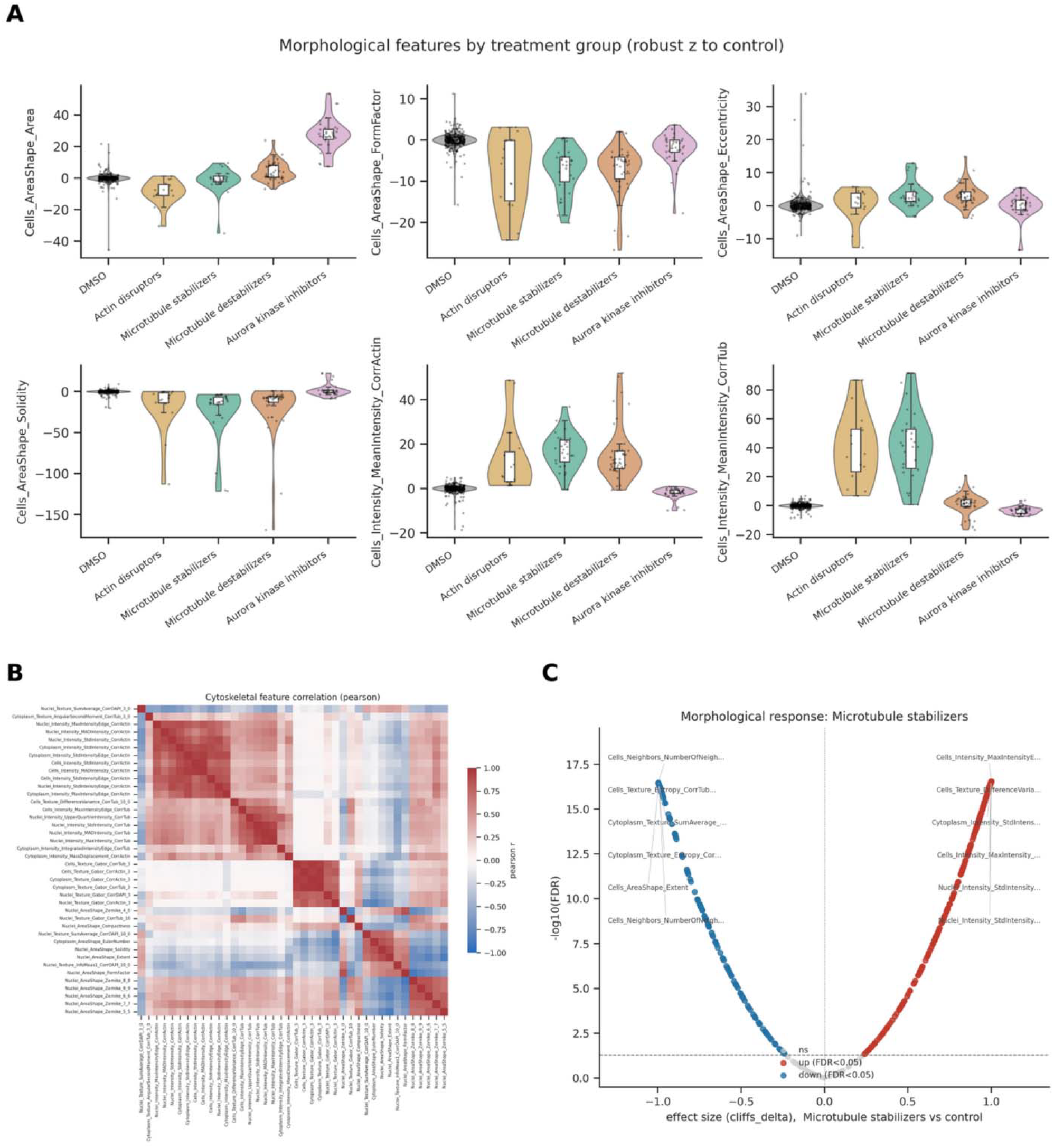
Representative violin plots for two texture features across selected MOA classes. Left panels show a nucleus texture feature that separates actin disruptors from DMSO controls (Mann–Whitney U, Cliff’s delta > 0.5, FDR < 0.01). Right panels show a cytoplasmic texture feature that distinguishes microtubule stabilizers. Each violin is overlaid with median and interquartile range.

## 4. Discussion

The per-feature test selection in MorphoStat avoids two errors common in morphological profiling studies. Using Student’s t-test across all features inflates false positives for the majority of features that fail normality. Defaulting to nonparametric tests for all features discards power for the features that happen to be normally distributed. The decision tree in Section 2.3 keeps both error rates under control at the cost of one additional test per feature, which is computationally negligible given the sample sizes typical of high-content screening runs.

MAD-based plate normalization was sufficient for BBBC021. The scaling factor 1.4826 is appropriate when control-well distributions are approximately normal. For datasets where DMSO variability is itself highly skewed, quantile normalization may work better, and a future version of MorphoStat will include that option.

MorphoStat currently accepts well-level summary profiles and does not model single-cell heterogeneity within a well. For experiments where bimodal or subpopulation-specific responses are expected, a single-cell analysis mode would be needed. That extension is planned for a future release.

MorphoStat is designed to complement imaging platforms rather than replace them. The intended workflow is: run CellProfiler or a compatible tool to produce a well-level CSV, then pass that file to MorphoStat. This boundary makes it straightforward to swap the upstream imaging software without modifying the downstream statistical analysis.

## 5. Conclusion

MorphoStat provides a complete analysis path from a CellProfiler output CSV to statistical results and figures, without manual scripting between steps. Automatic per-feature test selection, plate normalization against on-plate DMSO controls, and Benjamini–Hochberg correction reduce manual decisions and keep false-discovery risk low. Validation on the BBBC021 benchmark confirms that the pipeline recovers known pharmacological groups without manual feature curation. MorphoStat is available at https://github.com/Almunthir334/morphostat (DOI: 10.5281/zenodo.20354069) under the MIT license.

## Supporting information

TableS3_omnibus_across_classes

TableS2_pairwise_vs_control

## Acknowledgments

The authors thank the Broad Institute for making the BBBC021 dataset publicly available.

## Data Availability

The BBBC021 dataset is available from the Broad Bioimage Benchmark Collection at https://bbbc.broadinstitute.org/BBBC021. MorphoStat source code and analysis scripts are available at https://github.com/Almunthir334/morphostat (DOI: 10.5281/zenodo.20354069).

