## Supplementary_Material_Table1 for "MorphoStat: A Statistics-Aware Pipeline for Morphological Profiling Analysis"

*MorphoStat: automated, statistics-aware analysis of morphological profiling data*

*Contents: Supplementary Methods (S1–S5); Supplementary Table S1 (embedded); Supplementary Tables S2–S3 (provided as separate data files).*

### **Supplementary Methods**

**S1 Validation dataset preparation**

The validation data derive from the publicly available BBBC021 image set (Broad Bioimage Benchmark Collection). Per-well aggregated CellProfiler profiles were joined to the image metadata and to the curated compound mechanism-of-action (MOA) annotations on the plate and well keys; only wells with a defined MOA annotation were retained. The resulting analysis matrix contained 632 wells described by morphology features and metadata, spanning 13 MOA classes and anchored by 330 DMSO vehicle-control wells distributed across all plates. After cleaning (Section S3), 473 features were retained. One comma-separated input file per plate was written and passed to MorphoStat; the join is reproduced by the script provided in the software repository.

**S2 Batch-aware normalization**

Within each batch b (here, each plate), every feature x was transformed to a robust z-score relative to the negative control as x′ = (x − m_b_) / (1.4826 × MAD_b_), where m_b_ and MAD_b_ are the median and median absolute deviation of the control wells on plate b. The constant 1.4826 rescales the MAD to a standard-deviation–consistent estimate under normality. Plates with fewer control wells than a configurable minimum fall back to control statistics pooled across all plates. This centres each plate on its own vehicle and expresses all measurements in outlier-insensitive units comparable across plates.

**S3 Cleaning and distribution-aware statistical routing**

Feature columns were coerced to numeric type; columns with more than 50% missing values or with near-zero variance were removed, and remaining missing values were imputed by the column median. For each retained feature, normality was assessed by the Shapiro–Wilk test (large groups were randomly subsampled to keep the statistic well-behaved) and equality of variance by Levene’s test. Two-group comparisons with normal, homoscedastic data were routed to Student’s t-test; normal heteroscedastic data to Welch’s t-test; and non-normal data to the Mann–Whitney U test. Across more than two groups, one-way ANOVA and the Kruskal–Wallis test were the parametric and non-parametric routes. Effect sizes were the bias-corrected standardized mean difference (Hedges’ g) for parametric routes and the rank-based dominance statistic (Cliff’s δ, bounded in [−1, 1]) for non-parametric routes; η² and ε² were reported for ANOVA and Kruskal–Wallis. P-values for the entire family of feature-by-group comparisons were adjusted by the Benjamini–Hochberg procedure, and significance was declared at an adjusted false discovery rate below 0.05. On the BBBC021 profiles the control distributions were non-normal for essentially every feature, so all comparisons were routed to the non-parametric tests automatically.

**S4 FilamentSensor adapter**

For per-filament inputs, MorphoStat aggregates filament-level rows to per-image features: filament length and width are summarized by their means, orientation by the circular mean, and filament alignment by the nematic order parameter S = ⟨cos 2(θ − ⟨θ⟩)⟩, where θ is the filament angle. S ranges from 0 for isotropic (randomly oriented) filaments to 1 for perfectly aligned filaments. This adapter is provided for users’ own FilamentSensor exports; the validation in the main text used per-cell CellProfiler measurements.

**S5 Software environment and reproducibility**

MorphoStat is implemented in Python (≥3.9) using NumPy, pandas, SciPy, scikit-learn, statsmodels and Matplotlib/seaborn. Each run is deterministic and writes a machine-readable manifest (parameters, input fingerprint, software versions) and a Markdown report (cleaning audit, test-selection summary, counts of significant findings) in addition to the figures and statistics tables, so that an analysis is fully specified by its command and inputs. The complete validation reported here was produced by the single command:

morphostat run --input <plates> --group-col moa --control DMSO --batch-col Plate --source cellprofiler --normalization robust_z_to_control --impute median --alpha 0.05

### **Supplementary Tables**

**Table S1.** *Composition of the BBBC021 validation set and number of morphology features differing significantly from the DMSO control (Mann–Whitney U, Benjamini–Hochberg FDR < 0.05) for each mechanism-of-action class in the full 13-class screen (632 wells, 473 features).*

| **Mechanism-of-action class** | **Wells (n)** | **Significant features (of 473)** |
| --- | --- | --- |
| DMSO (negative control) | 330 | — |
| Microtubule destabilizers | 42 | 425 |
| Eg5 inhibitors | 36 | 423 |
| Microtubule stabilizers | 27 | 422 |
| Protein synthesis | 24 | 418 |
| Aurora kinase inhibitors | 36 | 405 |
| Epithelial | 22 | 397 |
| Cholesterol-lowering | 18 | 393 |
| Actin disruptors | 15 | 384 |
| Kinase inhibitors | 10 | 359 |
| DNA replication | 24 | 343 |
| DNA damage | 27 | 334 |
| Protein degradation | 21 | 316 |

**Table S2.** *Full per-feature pairwise statistics versus DMSO control for all mechanism-of-action classes (feature, group, selected test, raw P-value, Benjamini–Hochberg FDR, effect size and direction; 5676 rows). Provided as the data file TableS2_pairwise_vs_control.csv.*

**Table S3.** *Omnibus comparison across all 13 mechanism-of-action classes for each feature (Kruskal–Wallis test, ε² effect size, Benjamini–Hochberg FDR; 473 rows). Provided as the data file TableS3_omnibus_across_classes.csv.*
